# Chloramphenicol-mobilized *Bacillus subtilis* transiently expresses resistance to multiple antibiotics, including the glycopeptides phleomycin and bleomycin

**DOI:** 10.1101/2025.01.13.632840

**Authors:** Amelia Brave, Sandra LaBonte, Yongjin Liu, Morgan Powers, Evan De Ford, Paul D. Straight

## Abstract

Antibiotic resistance presents an urgent global crisis, exacerbated by antibiotic overuse. Understanding of the regulation of resistance genes within bacterial populations can inform strategies to prevent the spread of antibiotic resistance and reveal how antibiotics shape microbial communities. We identified upregulation of five antibiotic resistance loci in *Bacillus subtilis* colonies on solid growth media, following exposure to subinhibitory chloramphenicol concentrations. Notably, four resistance loci, *bmrCD, vmlR, tlrB,* and *ytbDE,* are regulated by transcription attenuation. Full expression depends upon antibiotic-induced ribosome stalling on upstream leader peptides, promoting transcription of the downstream gene. Here, we use luciferase reporter constructs fused to the 5’ regulatory region of each resistance gene to show differential spatiotemporal patterns of antibiotic resistance gene expression, revealing an intrinsic activation in addition to chloramphenicol induction in mobilized *B. subtilis* colonies. Because expression is under translational regulation, the data suggest natural translation pausing, in addition to antibiotic exposure, is an endogenous function that regulates these antibiotic resistance genes. While VmlR and TlrB have been previously characterized as conferring resistance to LS_A_Ps (lincosamides, streptogramin A, and pleuromutilin) and tylosin, respectively, antibiotics for BmrCD and YtbDE resistance complexes have not yet been identified. We demonstrate that although these resistance genes do not provide resistance to chloramphenicol, pre-exposure to chloramphenicol improved *B. subtilis* growth when cells were subsequently exposed to subinhibitory concentrations of respective antibiotics. We discovered that BmrCD confers resistance to the DNA-damaging glycopeptides, phleomycin and bleomycin, revealing that resistance arising from stalled ribosomes extends beyond drugs that target translation.

## INTRODUCTION

The rise of antibiotic resistance in pathogenic bacteria is increasingly becoming a global concern. While the origins of antibiotic resistance function vary, many are thought to protect bacteria from antimicrobial compounds produced by surrounding microorganisms in their natural environments (1, 2). Resistance mechanisms have likely arisen to provide microorganisms a competitive edge within their natural environment. These mechanisms are diverse, including efflux pumps, proteins protecting the ribosome, and proteins modifying antibiotics to prevent binding to their target (3) and have likely evolved to enhance bacterial fitness in diverse microbial communities, where organisms may encounter many antibiotics that target several essential processes (4).

*Bacillus subtilis* is a well-studied organism that serves as a model for many Gram-positive pathogens, including methicillin-resistant *Staphylococcus aureus* (MRSA). Therefore, understanding the regulatory mechanisms controlling antibiotic resistance gene expression in *B. subtilis* may provide valuable insight into resistance in clinically relevant bacteria. Previously, we used an interspecies competition model with *Streptomyces venezuelae* and *B. subtilis* on solid media and found that subinhibitory amounts of chloramphenicol (CHL) induced mobilization in *B. subtilis* (5), an example of hormesis, wherein inhibitory drugs are stimulatory at reduced concentrations (6, 7). Mobilization of a colony depends on sliding motility, driven by surfactin and exopolysaccharide production, and a substantial rewiring of metabolism (8–10). In addition to mobilization, subinhibitory CHL upregulates several antibiotic resistance genes, including *bmrCD*, *vmlR, tlrB*, *mdr,* and *ytbDE* (8, 11, 12) (Table 1). BmrCD is a multidrug efflux pump with a previously unknown substrate (11, 13), VmlR is an ATP-binding cassette type F (ABCF) ATPase providing resistance to LS_A_Ps (14, 15), TlrB is a methyltransferase conferring resistance to tylosin through methylation of 23S rRNA (12, 16), Mdr is a multidrug efflux pump predicted to provide resistance to fluoroquinolones (17), and YtbDE is a putative resistance complex (18). Consistent with prior studies, subinhibitory antibiotic exposure upregulates these unrelated resistance genes in *B. subtilis* (8, 11, 19).

**Table 1.** Antibiotic resistance genes upregulated upon exposure to subinhibitory concentrations of chloramphenicol.

|  | Function | Resistance | 6h<br>(log2fold)** |
| --- | --- | --- | --- |
| ***bmrCD*** | Heterodimer ABC transporter | ‡Phleomycin, bleomycin (glycopeptides) | 2.64,<br>2.26 |
| ***tlrB*** | 23S rRNA (guanine-N(1)-)-methyltransferase | Tylosin (macrolide) | 2.12 |
| ***vmIR*** | ARE ABCF ATPase | Streptogramins (macrolide), pleuromutilins, lincomycin (glycopeptide) | 2.27 |
| ***mdr*** | multidrug-efflux transporter | Puromycin* (aminonucleoside),<br>nerfloxacin*, tosufloxacin* (fluoroquinolone) | 2.03 |
| ***ytbDE*** | D - similar to antibiotic resistance<br>E - putative aldo/keto reductase | Unknown | 2.56,<br>2.48 |

‡Identified in this study.
\*Predicted based on structure and sequence similarity but not experimentally demonstrated
\*\*log2fold change in transcript abundance extracted from *B. subtilis* colonies exposed to subinhibitory (1 µM) chloramphenicol compared to untreated wild type *B. subtilis* colonies from transcriptomics study in Liu et al. 2024 (8)

Of the five resistance loci showing increased expression in mobilized colonies, four (*bmrCD, vmlR, tlrB,* and *ytbDE*), are regulated by common transcription attenuation mechanisms (11, 12, 20, 21). Each contains an upstream open reading frame (uORF) encoding a leader peptide whose nascent mRNA secondary structure controls downstream gene expression. Intrinsic formation of an anti-antiterminator stem loop and a terminator stem promotes the dissociation of RNA polymerase upstream of the antibiotic resistance ORF due to NusG-dependent pausing, thus preventing transcription. Antibiotic-induced ribosome stalling allows time for the formation of an antiterminator hairpin, prevents formation of a terminator stem loop, and thus inhibits expression of the downstream gene (Figure 1A, S1A, S1B) (11, 12, 15).

**Figure 1.**
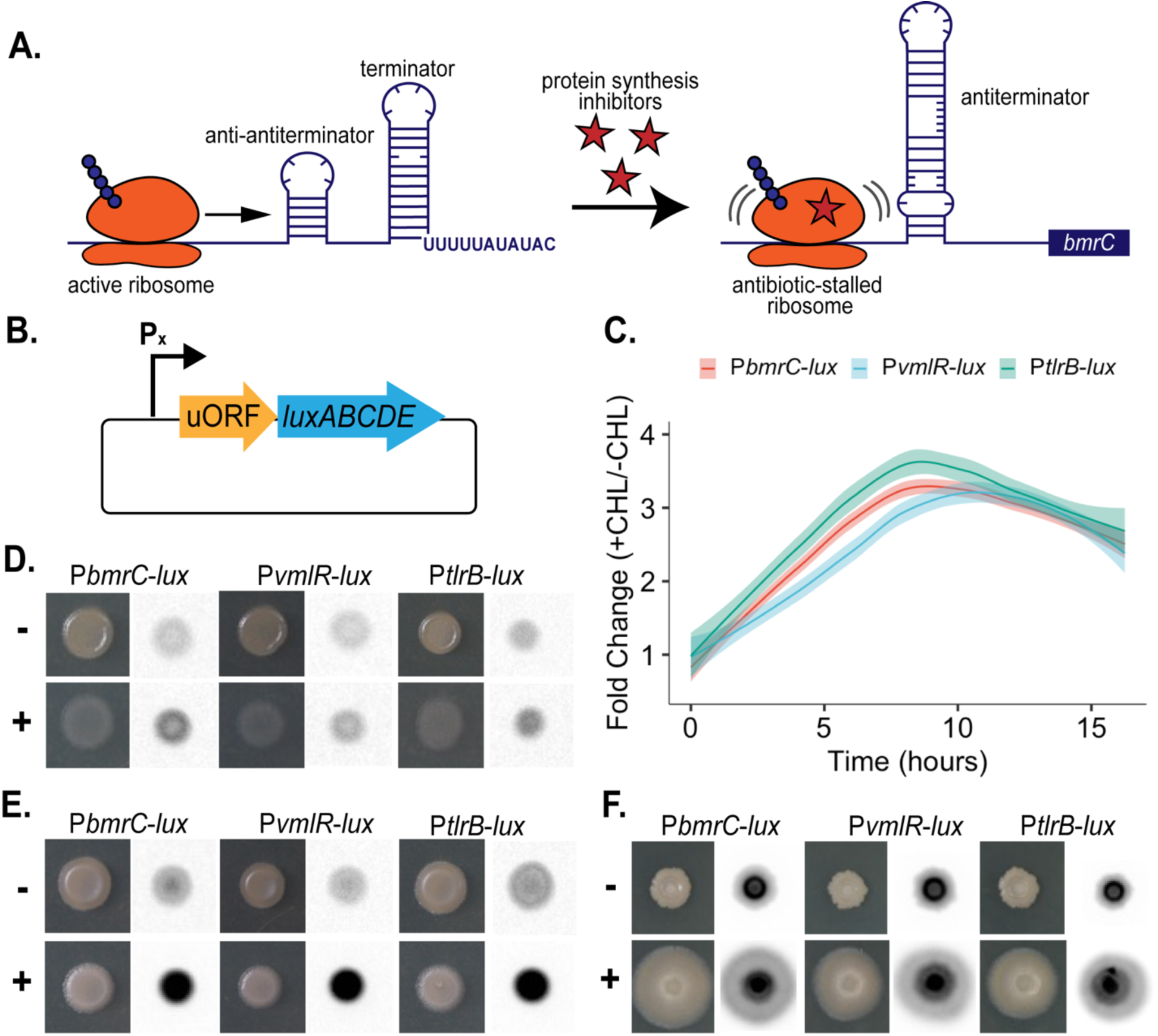
Transient and spatiotemporal expression of the antibiotic resistance factors BmrCD, VmlR, and TlrB in the presence of chloramphenicol. **A.** Overview of the leader peptide transcription attenuation regulatory mechanism, using *bmrCD* as an example. In the absence of ribosome stalling, the nascent transcript encoding the uORF forms anti-antiterminator and terminator hairpins. RNA polymerase dissociates upstream of the resistance ORF. Antibiotic-induced ribosome stalling allows for the formation of an antiterminator hairpin, which allows transcription of the downstream resistance gene. **B.** Reporter construct design. We used the native promoter (P_x_) and leader peptide gene for each resistance factor (*bmrCD, vmlR, tlrB,* and *ytbDE*) fused to the bacterial luciferase cassette. **C.** The fold change in relative light units (RLU) of P*bmrC-lux,* P*vmlR-lux,* and *PtlrB-lux* exposed to 1 μM CHL compared to non-exposed grown over 16 hours. The curves were normalized to a luciferase construct lacking a promoter. **D-F.** (**Note:** The chemiluminescence images are inverted. The luminescence signal appears gray to black with increasing intensity) P*bmrC-lux,* P*vmlR-lux,* and *PtlrB-lux* spotted on agar plates with 1 μM CHL (+) or without CHL (-). Pictures were taken with phase contrast (left) and chemiluminescence (right) at time points **D.** 6 hours **E.** 9 hours and **F.** 24 hours. For scale, the 24 h colonies are ∼0.5 cm in diameter, except in F., +CHL, where the colonies have mobilized to approximately twice the initial diameter.

*B. subtilis* forms specialized subpopulations that exhibit different spatiotemporal gene expression patterns dependent on growth conditions (8, 22, 23). Therefore, to assess their expression in a dynamic surface-mobilized population, we investigated the spatiotemporal expression patterns of these resistance genes following ribosome stalling due to subinhibitory CHL exposure. We use engineered reporter strains containing fusions of the 5’ regulatory regions containing the promoter and uORF to bacterial luciferase genes. Although CHL induces expression of all four reporters, we found no evidence of CHL resistance nor influence on sliding colony morphology provided by any of the resistance genes. Pre-exposure to CHL, however, improved growth when cells were subsequently exposed to the corresponding antibiotic for each resistance function. Two additional observations indicate that transient multidrug resistance is activated during pausing of translation. First, we found that on solid media without CHL, the reporter constructs were transiently expressed, revealing an intrinsic pause in translation leading to upregulation of multiple drug resistance functions. Second, we discovered that the BmrCD efflux pump provides resistance to the glycopeptides, phleomycin and bleomycin, that target DNA, not protein synthesis. These results suggest a model wherein *B. subtilis* may gain a competitive advantage by entraining antibiotic resistance to translation rates. Antibiotic exposure promotes intrinsic resistance to multiple antibiotics, which in concert with mobilization, elevates fitness in the presence of antibiotic-producing competitors.

## RESULTS

### Spatiotemporal patterning of resistance genes expression in *B. subtilis* colonies exposed to CHL

Previously, we identified approximately 800 genes significantly (>log2fold change, padj < 0.05) differentially expressed in *B. subtilis* NCBI 3610 sliding colonies triggered by subinhibitory CHL exposure (8). From this dataset, we identified five characterized or putative antibiotic resistance loci upregulated at both 6 hours and 24 hours post CHL exposure: *bmrCD, vmlR, tlrB, ytbDE,* and *mdr*. Of these, all but *mdr* are regulated by transcriptional attenuation mediated by antibiotic-induced ribosome stalling on a 5’uORF (11, 12, 15, 20) (Figure 1A, S1A, S1B). We sought to monitor temporal differences in the expression of these four genes, regulated by a similar mechanism in the presence of CHL, and to identify subpopulations in a mobilized colony expressing these resistance factors. To monitor expression, we constructed reporter strains by fusing the native promoter and regulatory 5’uORFs for each respective gene to luciferase (Figure 1B). We first monitored luciferase expression of these reporter strains in liquid cultures to determine if CHL triggers expression of these resistance genes, since our prior transcriptomics data was from colonies on solid media. We observed similar expression profiles in the characterized resistance genes, *bmrCD, vmlR,* and *tlrB.* Compared to cultures lacking CHL, liquid cultures with 1 μM CHL demonstrated an upregulation of resistance gene transcriptional activity, peaking after 8 to 12 hours of growth and then decreasing (Figure 1C). We additionally found low to no uninduced expression of the resistance genes compared to a control construct lacking a promoter (Figure S2), which is consistent with prior studies (11, 15, 21). Because liquid cultures are homogenous compared to the diverse subpopulations present in a colony on solid media, we then monitored spatiotemporal expression of the reporter strains over a 48-hour period (Figures 1D-1F). Like in liquid cultures, we observed transcriptional activity of the resistance genes upregulated by the presence of CHL. At 6 and 9 hours of growth, we observed increasing expression throughout the colony over time (Figure 1D, 1E). At 24 hours, we observed an actively sliding colony with expression primarily present in the center of the colony (Figure 1F). After 48 hours of growth, this expression has moved from the center of the colony to the subpopulation at the edge of the outward expanding colony (Figure S3). Surprisingly, we also observed transcriptional activity at 24 hours in colonies grown without antibiotic (Figure 1F). The expression differed in intensity and pattern from colonies exposed to CHL at 9 hours (Figure 1E) and 24 hours yet revealed substantial induction in all three reporter strains. The highest intensity was seen in a ring around the core of the colony in contrast to the center subpopulation transcriptional activity induced by CHL (Figure 1F). At 48 hours, like in CHL-exposed colonies, the transcriptional activity shifted to the outer the edge of the non-mobilized colony (Figure S3). The putative resistance genes, *ytbDE*, exhibited a different spatiotemporal pattern compared to the others (Figure S4). We observed promoter activity for *ytbDE* in both CHL exposed and untreated colonies, highest at 6 hours of growth, and diminishing at later times. Although all four resistance functions use a 5’uORF regulatory structure, the different spatiotemporal activity pattern for *ytbDE* indicates an additional regulatory function controlling its expression. The *ytbDE* uORF is unique among the four in that an oppositely directed ORF is overlapping in the genome, which may influence its expression control (18).

### Deletion of antibiotic resistance genes does not affect *B. subtilis* sensitivity to CHL

VmlR and TlrB are characterized resistance factors conferring resistance to LS_A_Ps and tylosin respectively. An antibiotic substrate for the multidrug ABC transporter BmrCD has not been previously identified, but it is active as an efflux pump that will transport fluorescent molecules in a vesicle assay (24). Mdr is a multidrug efflux pump predicted to provide resistance to fluoroquinolones (17). YtbDE has not yet been characterized. Although CHL exposure triggers transcriptional activity of these resistance factors, there is no evidence these genes provide resistance to CHL. Therefore, we tested if these resistance factors provide any resistance to CHL exposure or influence the hormetic stimulatory effect of subinhibitory CHL exposure. To test this, we deleted each of the five resistance genes and observed the response of these mutant strains to no CHL, subinhibitory CHL (1 μM), and inhibitory CHL (8 μM). We saw no difference in inhibitory concentration or colony morphology in response to stimulatory concentrations of CHL (Figure S5). To determine if these factors work synergistically to provide resistance to CHL, we deleted all 5 resistance genes and tested the strain (Δ5) with varying CHL concentrations. We again found no difference in inhibitory concentration or response to subinhibitory CHL (Figure 2). Therefore, although induced in the presence of CHL, these resistance genes do not mediate the response of *B. subtilis* to CHL.

**Figure 2.**
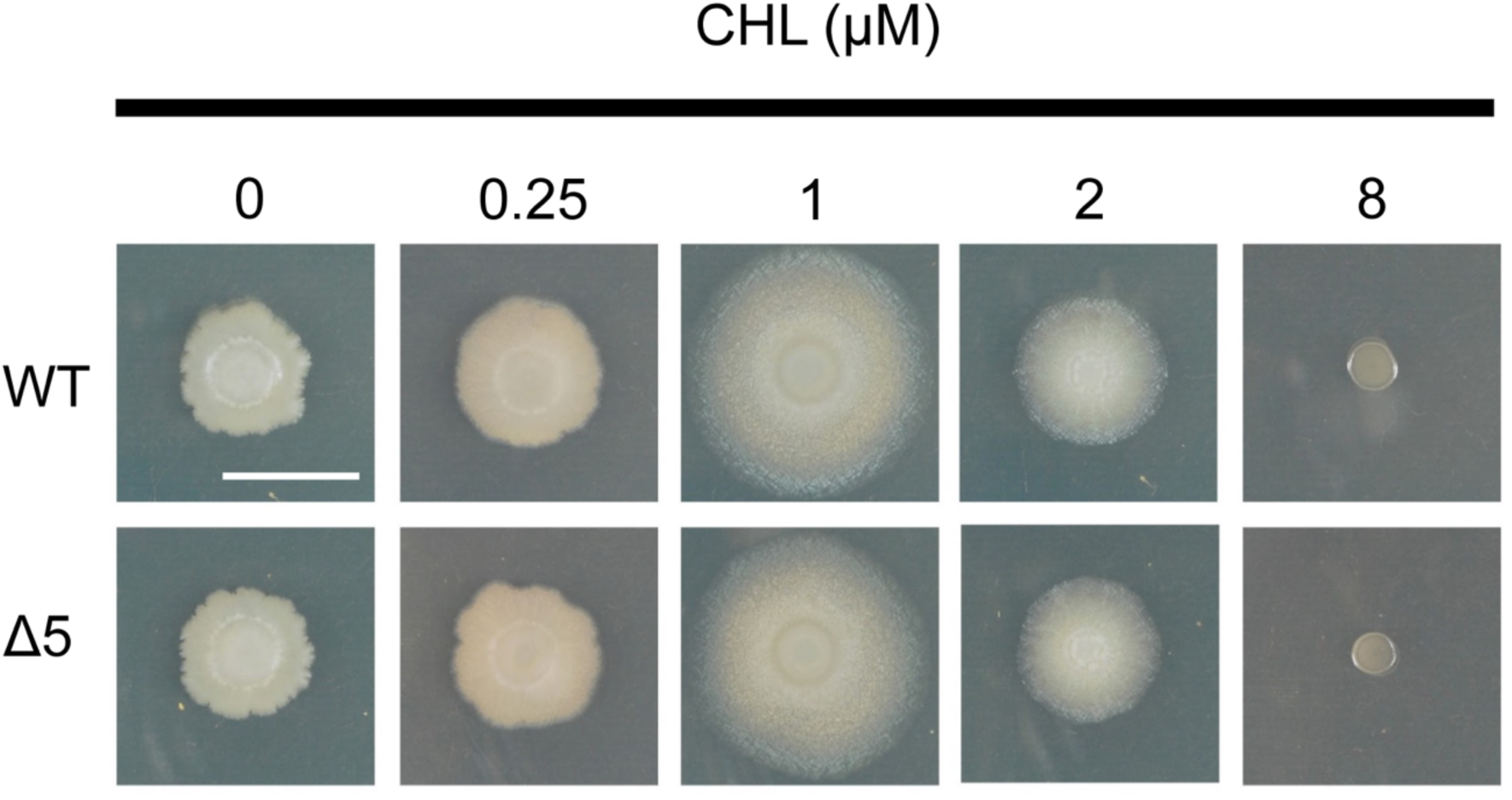
The antibiotic resistance factors upregulated in the presence of chloramphenicol do not provide resistance nor do they impact *B. subtilis* motility. Five resistance elements (*bmrCD*, *vmlR*, *tlrB*, *mdr* and *ytbDE*) that respond to CHL at 6 h were deleted (refer to as Δ5 strain). Wild type (WT) and Δ5 strains were plated on GYM7 agar with a range of subinhibitory chloramphenicol (0-8 μM). Pictures were taken at 24 h. Bar, 1 cm.

### Pre-exposure to CHL enhances *B. subtilis* growth in the presence of lincomycin and tylosin

Previously it was shown pre-exposure to tylosin promotes resistance through *tlrB* expression (12). We hypothesized that pre-exposure to CHL would promote resistance to tylosin and lincomycin through expression of *tlrB* and *vmlR* respectively. To test this, we pre-exposed *B. subtilis* cultures to 1 μM CHL and compared growth over 18 hours when subsequently exposed to CHL, lincomycin, or tylosin. We performed two-fold serial dilutions (from 32 to 0.03125 μg/mL) of each antibiotic and compared growth of the non-exposed and pre-exposed cultures to identify a change in minimum inhibitory concentration (MIC). We did not observe a significant change in MIC between the non-exposed and pre-exposed cultures in subsequent antibiotic treatment. This result is likely due to tylosin and lincomycin each inducing their specific resistance genes. We did, however, observe a difference in growth of the pre-exposed cultures compared to the controls when exposed to increasing concentrations of drug (Figure S6). We reasoned that CHL exposure would position the colony to better tolerate early exposure to corresponding antibiotics by priming specific resistances. To quantify this differential growth, we calculated IC_50_ values, the concentration of half maximal growth inhibition, after 4 hours of growth, which corresponds to late log phase of a wild-type culture grown in isolation under these conditions. We plotted the OD_600_ values at each concentration for pre-exposed and non-exposed (control) cultures, each normalized to cultures subsequently grown in the absence of antibiotic, to control for any growth differences due to CHL pre-exposure. We then used a sigmoidal fit to calculate IC_50_ values for pre-exposed and control cultures and compared these values. For cultures grown in CHL, we found no significant change in IC_50_ value following CHL pre-exposure (Figure 3A). This is likely because there are no known genes in *B. subtilis* for CHL resistance. There was, however, a statistically significant change in IC_50_ value of cultures pre-exposed to CHL in lincomycin-exposed (Figure 3B) and tylosin-exposed (Figure 3C) cultures. We concluded that CHL pre-exposure provides a resistance benefit from early expression of the resistance factors *vmlR* and *tlrB*. Although this likely arises from priming resistance to lincomycin and tylosin, it may in part arise alternatively from an undefined role in global metabolic control, leading to faster growth, which could explain the accelerated growth in cultures pre-exposed to CHL and subsequently grown in isolation (Figure S6). However, this is unlikely because the same benefit is absent when cells are subsequently cultured with CHL instead of lincomycin or tylosin (Figure 3A).

**Figure 3.**
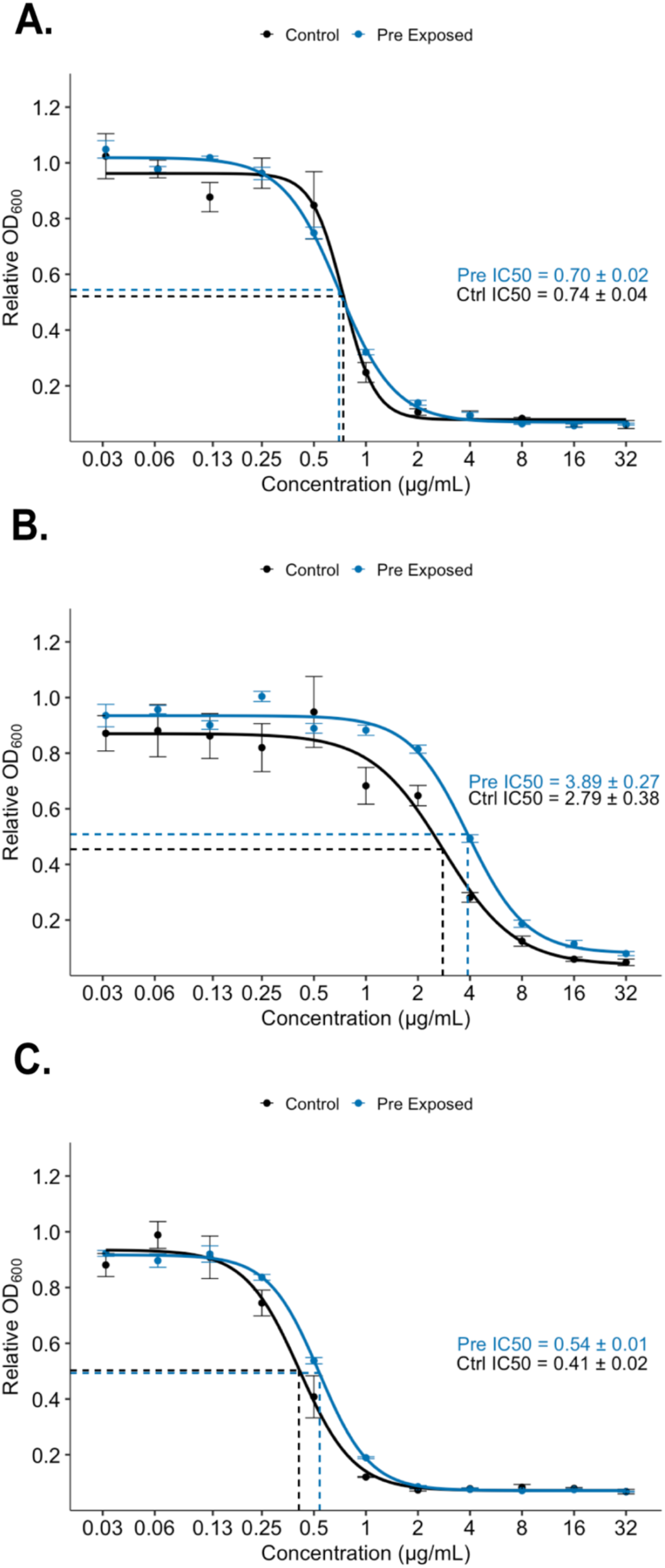
Pre-exposure to chloramphenicol improves *B. subtilis* tolerance to subinhibitory concentrations of lincomycin and tylosin. *B. subtilis* overnight cultures were diluted to OD_600_ = 0.08 and grown with (pre-exposed) or without (ctrl) 1 μM CHL for 2.5 hours. For each sample well, 5 × 10^5^ cells were inoculated and grown in a 2-fold dilution series of **A.** CHL, **B.** lincomycin, or **C.** tylosin. The relative OD_600_ value after 4 hours of growth was plotted over concentration and fit with a sigmoidal curve. Each data point is relative to the OD_600_ of pre-exposed or control respectively grown without antibiotic. IC_50_ values are the concentration at half the maximum OD_600_.

### CHL treatment leads to increased phleomycin and bleomycin resistance through induction of BmrCD expression

We next questioned whether the four resistance functions, *bmrCD*, *ytbDE, vmlR,* and *tlrB,* may share a transcription attenuation regulatory mechanism because they provide resistance against protein synthesis inhibitors. Therefore, we sought to identify antibiotics for which BmrCD and YtbDE provide resistance. To identify a substrate antibiotic, we compared sensitivity of Δ*bmrCD* and Δ*ytbDE* strains with wild-type *B. subtilis* sensitivity to a panel of antibiotics. In the case of Δ*ytbDE*, we did not observe elevated sensitivity to any of the antibiotics tested in this study (Table S1). The Δ*bmrCD* strain, while not showing increased sensitivity to any protein synthesis inhibitors, instead displayed increased sensitivity to phleomycin compared to wild type *B. subtilis* (Figure 4A). Following addition of 1µM CHL to the agar medium, we observed resistance to phleomycin in the wild-type strain but not the Δ*bmrCD* mutant strain, suggesting BmrCD confers resistance to phleomycin (Figure 4A). Because phleomycin belongs to a large class of diverse glycopeptides, we selected two additional representative glycopeptides to test for resistance, bleomycin and vancomycin. The Δ*bmrCD* strain revealed sensitivity to both glycopeptides, although bleomycin sensitivity was increased in the Δ*bmrCD* strain compared to wild type (Figure 4A). Addition of 1µM CHL enhanced the bleomycin resistance in the wild-type strain, whereas sensitivity was unaffected in Δ*bmrCD* and in either strain when exposed to vancomycin. Phleomycin and bleomycin are structurally similar glycopeptide antibiotics that cause oxidative DNA damage and lead to single and double strand breaks (25). Vancomycin is a structurally different glycopeptide that targets the cell wall (26). Therefore, the results indicate that BmrCD provides selective resistance for the phleomycin/bleomycin subtype of glycopeptides with cytoplasmic targets.

**Figure 4.**
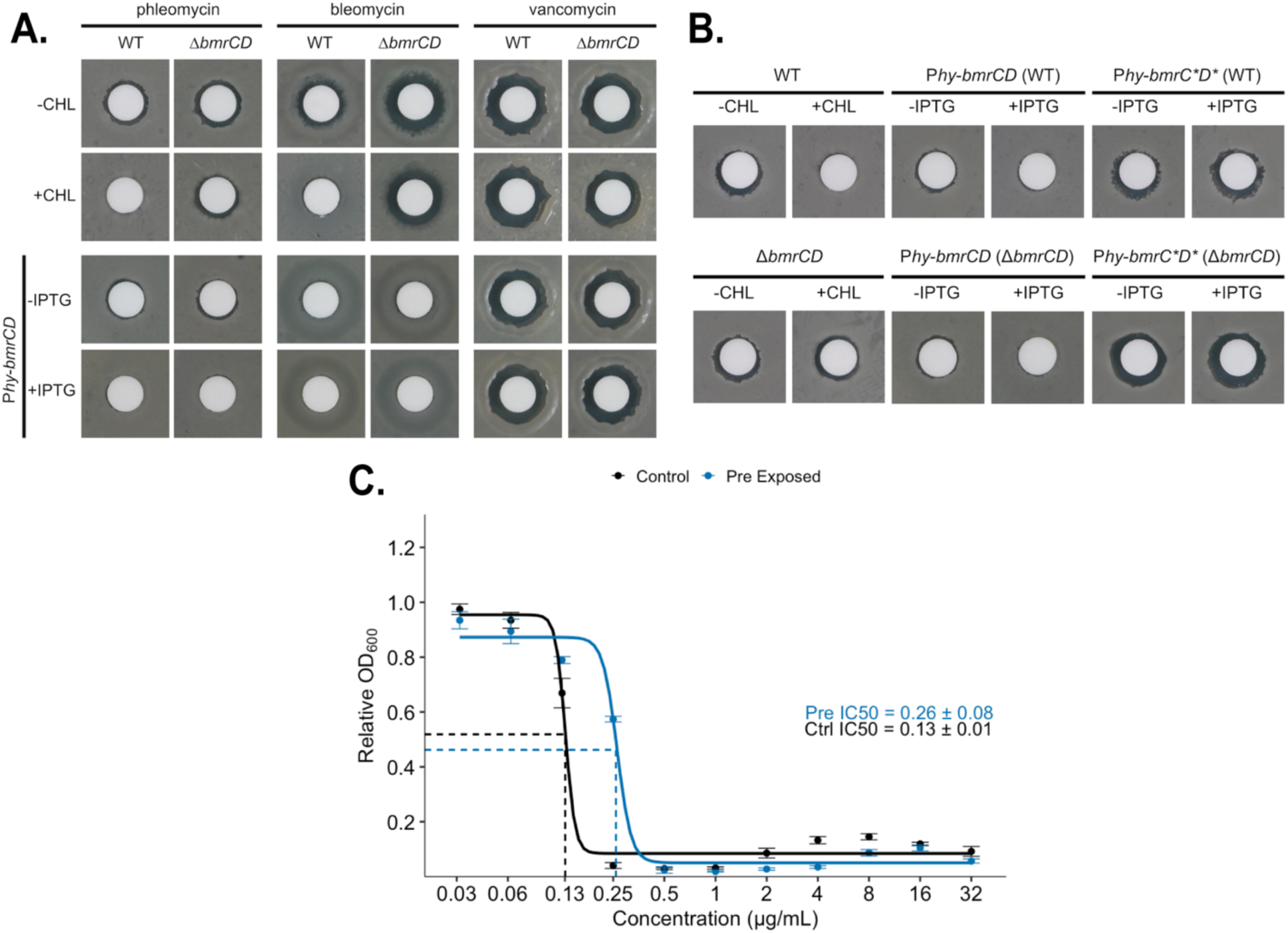
BmrCD confers resistances to the glycopeptides phleomycin and bleomycin. **A.** Left: WT and Δ*bmrCD* strains (OD600=1.0, 100 μl) were spread on the GYM7 agar medium in the absence (-) and presence of 1 μM CHL. Then, 10 μl of 125 μg/ml phleomycin was added onto the paper disc. Middle: Complementation of Δ*bmrCD* with *bmrCD* under the control IPTG-inducible promoter (*hyperspank*) restored the phleomycin resistance in the presence of 0.5 mM IPTG, with an observation of reduced inhibitory zone. There is some leaky expression for this promoter. Right: point mutations introduced to Walker A motif of BmrC and BmrD in both WT and Δ*bmrCD* strains abolished the BmrCD function as a phleomycin transporter, with an observation of inhibitory zone similar to the group without IPTG. Pictures were taken at 24 h. Diameter of paper disc, 6 mm. **B.** WT and Δ*bmrCD* strains (OD600=1.0, 100 μl) (top) and WT and Δ*bmrCD* strains with P*hy-bmrCD* inserted at the *amyE* locus (OD600=1.0, 100 μl) (bottom) were spread on the GYM7 agar medium in the absence (-) and presence of 1 μM CHL. Then, 10 μl of 125 μg/mL phleomycin (left), 10 μl of 1 mg/mL bleomycin (middle) and 10 μl of 25 μg/ml vancomycin (right) were added onto each individual paper disc. Pictures were taken at 24 h. Diameter of paper disc, 6 mm. **C.** *B. subtilis* overnight cultures were diluted to 0.08 and grown with (Pre) or without (Ctrl) 1 μM CHL for 2.5 hours. 5 × 10^5^ cells from these subcultures were grown in a 2-fold dilution series of phleomycin. The relative OD_600_ value after 4 hours of growth was plotted over concentration and fit with a sigmoidal curve. Each data point is relative to the OD_600_ of pre-exposed or control, respectively, grown without antibiotic. IC_50_ values are the concentration at half the maximum OD_600_.

To confirm the function of BmrCD in phleomycin and bleomycin resistance, we complemented the loss-of-function by expressing *bmrCD* under the IPTG-inducible promoter P*hyperspank* in the Δ*bmrCD* strain. This resistance to phleomycin and bleomycin was restored in the complemented mutant, even in the absence of IPTG, which we attributed to leaky expression under the strong promoter regulation (Figure 4A). As expected, resistance to vancomycin was unaffected. Therefore, to determine whether resistance is dependent upon active BmrCD, we next engineered a strain expressing alleles of *bmrCD* with point mutations in the Walker A motifs of each gene (BmrC K377A and BmrD K469A), again under the P*hyperspank* promoter (Figure S7), preventing nucleotide binding in BmrCD and eliminating its ability to transport substrates (27). This mutant was incapable of providing phleomycin or bleomycin resistance in the presence or absence of IPTG, demonstrating that active BmrCD conveys resistance to these DNA-damaging glycopeptide antibiotics (Figure 4B).

Contrary to our hypothesis that the CHL-induced genes selectively protect protein synthesis, the observed resistance to DNA-damaging antibiotics reveals that CHL exposure elevates a broader range of intrinsic antibiotic resistance. We next sought to determine whether CHL-induction of *bmrCD* expression provides a quantitative increase in IC_50_ as observed for lincomycin and tylosin. We monitored growth and determined IC_50_ values identically as done previously (Figure 3), comparing growth in the presence and absence of phleomycin following pre-exposure to subinhibitory CHL. Comparing growth at 4 hours (late log phase), we measured an increase in IC_50_ value following pre-exposure to CHL for induction of *bmrCD* expression (Figure 4C). Unlike with lincomycin and tylosin, we also observed a transient drop in OD_600_ of the control cultures with 0.25 μg/mL phleomycin and no growth with higher concentrations (Figure S6D). At 0.25 μg/mL phleomycin, the CHL pre-exposed colonies grew to an OD_600_ roughly equivalent to the no phleomycin control, dropping again after 10 hours of growth, possibly due to cell lysis. The growth curve difference of the phleomycin-exposed cultures may be attributed to the SOS response in *B. subtilis* providing some resistance to phleomycin (28). Because phleomycin targets DNA and not protein synthesis, exposure to phleomycin should not induce expression of the *bmrCD* genes. Using the luciferase assay for expression, we found the presence of phleomycin does not induce *bmrCD*, consistent with it not inducing ribosome stalling (Figure S8).

## DISCUSSION

Antibiotics are one form of complex chemical traffic between species within microbial communities. A persistent question is how antibiotics function in communities and whether they reach clinically inhibitory concentrations. A complementary question is how antibiotic resistance may operate in microbial communities. Our lab previously identified that CHL produced by *S. venezuelae*, although inhibitory at higher concentrations, stimulates sliding motility in *B. subtilis* colonies at subinhibitory concentrations (5). In these mobilized colonies, *B. subtilis* upregulates several antibiotic resistance genes in response to the external stressor (8). Of the five antibiotic resistance loci we identified as upregulated in response to CHL, we found no evidence that any provide resistance to CHL. Here, we studied four of these resistance loci, commonly regulated through a terminator attenuator mechanism that is responsive to translation efficiency: *bmrCD, vmlR, tlrB,* and *ytbDE*. We found evidence of a cross-resistance phenomenon, where exposure to one antibiotic, CHL, triggers the expression of resistance genes specific to unrelated antibiotics. Pre-exposure to CHL improves the growth of *B. subtilis* grown in the presence of lincomycin and tylosin by triggering the expression of *vmlR* and *tlrB*. Lincomycin and tylosin are protein synthesis inhibitors like CHL, which suggested that any form of translation stress will stimulate multidrug resistance due to the physiological impact on ribosome activity, as opposed to a one-to-one correspondence between a specific antibiotic and resistance function.

Expression of *bmrCD, vmlR,* and *tlrB* occurred in similar spatiotemporal profiles when cultured on agar media. We saw consistent strong expression throughout the colony at early growth stages in the presence of subinhibitory CHL. In an actively sliding colony we observed strong expression in the center of the colony, and over time the pattern shifts to expression at the colony edge. The observed pattern suggests that specialized subpopulations transiently express resistance in the presence of translation inhibitors, while other subpopulations likely play different roles in driving colony mobilization. Unexpectedly, we observed transient expression of all resistance genes in the absence of CHL on solid media, with *bmrCD, vmlR,* and *tlrB* exhibiting similar spatiotemporal expression profiles. Previously, it was shown *B. subtilis* evolved a tight regulation system, preventing unnecessary expression of these energetically costly resistance genes. These resistance genes share a common regulatory mechanism involving antibiotic-dependent ribosome stalling mediating transcription attenuation (11, 12, 15). NusG-dependent pausing provides time for leader peptide translation that is essential for coupling ribosome position to transcription termination. Leaky expression of *vmlR* and *tlrB* are further controlled by a translation attenuation mechanism where transcripts evading termination are silenced by Shine-Dalgarno-sequestering hairpins (Figure S1A, S1B) (12, 15). In addition, *vmlR* expression is tightly regulated by NusA-stimulated transcriptional pausing and (p)ppGpp-mediated signaling (15). This combination of regulatory mechanisms is thought to have evolved to minimize the fitness cost associated with leaky expression in the absence of antibiotic pressure. However, our data reveal that within surface-grown colonies, the resistance genes are expressed without antibiotic exposure, suggesting either an endogenous pause in protein synthesis during growth that triggers expression, or alternatively, a regulatory control with an undefined mechanism.

Our initial hypothesis was that the common regulation of the four resistance genes is a general mechanism for resistance to translation inhibitors. However, we identified two substrates for the ABC transporter, BmrCD, as glycopeptide antibiotics, phleomycin and bleomycin. Unlike lincomycin (VmlR) and tylosin (TlrB), phleomycin and bleomycin do not target translation, but instead cause oxidative stress inducing single and double strand DNA breaks (25). Accordingly, phleomycin does not induce *bmrCD* expression, which is regulated through transcriptional attenuation trigged by ribosome stalling. Therefore, our data show that in the absence of a protein synthesis inhibitor, *B. subtilis* produces BmrCD transiently by sensing endogenous ribosome stalling, which may then provide resistance to DNA oxidative stress. Exposure to CHL amplifies and sustains BmrCD, revealing resistance to phleomycin and bleomycin. A structurally and functionally different glycopeptide, vancomycin, is not a substrate for BmrCD. Possibly, an unidentified translation inhibitor that shares common structural features with phleomycin and bleomycin could specify BmrCD transport, serving as a canonical trigger for antibiotic-specific resistance. Nevertheless, within a microbiome surrounded by various microorganisms producing diverse antibiotics, a common regulatory mechanism for expression of resistance to diverse antibiotic challenges may be advantageous.

We were unable to identify a substrate antibiotic for YtbDE. We suspect it provides resistance to an antibiotic not tested in this study. Because *mdr* is not regulated in the same way as the other four resistance genes, we did not assess its spatiotemporal expression or identify a substrate antibiotic, though Mdr is a multidrug efflux pump predicted to provide resistance to puromycin, norfloxacin, and tosufloxacin (17). Because *mdr* is not regulated in the same way but is upregulated in the presence of CHL, this suggests cross-resistance mechanisms in *B. subtilis* are not limited to the transcription attenuation regulatory mechanism evaluated in this study. Recently, Meirelles et al. also reported that phenazine produced from *P. aeruginosa* increased the tolerance to ciprofloxacin and levofloxacin in *P. aeruginosa* by upregulating the expression of *metGHI-opmD* efflux system (29). These data suggest that cross-resistance induced by different chemical stimuli may be a common phenomenon, which may provide a protective effect for bacteria against competitors producing diverse chemicals.

Physiological adaptations provide *B. subtilis* a complementary means to compete with surrounding microbes. In this study we have demonstrated a cross-resistance expression system where CHL triggers the expression of several unrelated antibiotic resistance genes. CHL exposure causes ribosome stalling, which allows time for a structural change in the nascent transcript encoding the regulatory leader peptide. Though ordinarily containing anti-antiterminator and terminator hairpins, the stalled ribosomes promote formation of an antiterminator structure. The downstream resistance gene is then expressed (Figure 5). We have shown the expression of these genes is tied to changes in growth of bacterial cells. These changes have been observed in heterologous populations of bacteria that show differential patterns of resistance gene expression, suggesting that during sensitive transitions in growth or exposure to subinhibitory antibiotics, bacteria encode mechanisms to prime intrinsic resistance function and boost competitive fitness. In summary, our study demonstrates *B. subtilis* responds to subinhibitory CHL exposure by inducing cross-resistance mechanisms to unrelated antibiotics, a phenomenon likely advantageous in diverse microbial environments. These findings reveal resistance gene expression is intricately linked to colony physiology and may serve broader endogenous roles, highlighting the complexity and adaptability of bacterial fitness in competitive natural environments.

**Figure 5.**
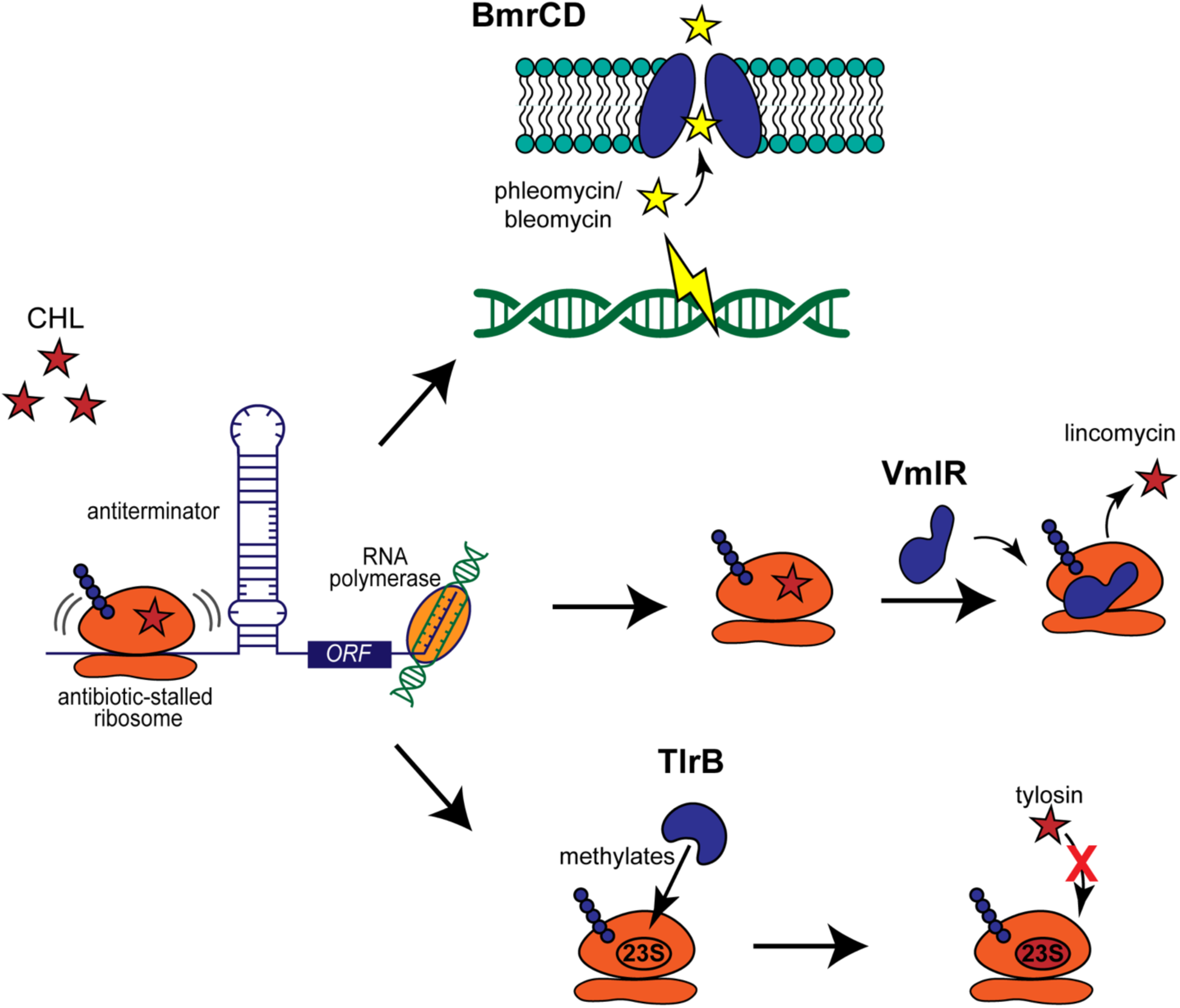
Model illustrating how expression of several antibiotic resistance genes is triggered by CHL exposure. CHL stalls the ribosome, preventing formation of a terminator stem loop. The downstream resistance genes regulated by this transcription attenuation mechanism, *bmrCD, vmlR,* and *tlrB,* are transcribed and translated. Two of these genes, *vmlR* and *tlrB*, confer resistance to protein synthesis inhibitors, lincomycin and tylosin, respectively, but not CHL. VmlR binds to the ribosome displaces L_s_AP antibiotics inhibiting translation. TlrB methylates the 23S rRNA to prevent tylosin binding to the ribosome. BmrCD expression is also induced by CHL exposure but instead offers resistance to glycopeptides phleomycin and bleomycin, both of which induce oxidative stress and double and single strand breaks in DNA. BmrCD is a multidrug efflux pump that transports phleomycin and bleomycin out of the cell.

## MATERIALS AND METHODS

### Strains, primers, and media

The strains of *Bacillus subtilis* used in this study are listed in **Table S2**. *Bacillus subtilis* mutant strains in 168 or PY79 background were transduced to NCIB3610 by SPP1 phage transduction (30). The primers are listed in **Table S3**. *Bacillus subtilis* strains were cultured at 37℃ in lysogeny broth (LB) and were inoculated onto GYM7 plates (0.4% [w/v] D-glucose, 0.4% [w/v] yeast extract, 1.0% [w/v] malt extract, pH7.0) with 1.5% [w/v] agar when grown to an OD600 of 1.0.

### Construction of luciferase reporter strains

To construct luciferase reporter strains, we used primers for each region upstream of the antibiotic resistance ORFs (listed in **Table S3**) to amplify the target leader peptide uORF and promoter from *B. subtilis* NCIB 3610 genomic DNA, primers *lux*-For and *lux*-Rev to amplify the *luxABCDE* fragment from pBS3Klux, primers *amyE*-back-For and *amyE*-front-Rev to amplify the plasmid backbone (including the origin site, ampicillin resistance cassette, and front and back partial *amyE* fragments) from pDR111, and primers *kan*-For and *kan*-Rev to amplify the kanamycin resistance cassette from pDG780 (31). These 4 fragments were assembled to a functional plasmid by Gibson assembly (32). The plasmid was transformed to *B. subtilis* PY79 wild type and then the construct containing the target 5’ region, *lux* operon, and kanamycin resistance cassette was inserted into the *amyE* locus. The inserted construct was verified by PCR. Once confirmed, the construct was moved to PDS0066 (*B. subtilis* NCIB 3610 wild type) using SPP1 phage transduction.

### Luciferase expression assay in liquid cultures

Strains were inoculated in LB and grown overnight at 37°C. The overnight was then diluted and grown to an OD600 of 1. Cultures were then diluted to an OD600 of 0.08 using liquid LB as diluent and 200 μL was dispensed in 96-well plates. Each luciferase strain was monitored in the presence of 1 μM CHL and without. Plates were incubated in an Agilent BioTek Synergy H1 microplate reader at 30 °C with continuous orbital shaking. Luminescence and OD_600_ was monitored every 15 minutes for 16 hours. All assays were performed with four biological replicates per condition. Luciferase expression was normalized to a luciferase strain lacking a promoter.

### Luciferase expression assays on solid media

*B. subtilis* cells grown in 4 mL LB broth were diluted to an OD600 of 0.08. When grown to an OD600 of 1.0, 1.5 μL of *B. subtilis* cells were spotted on the GYM7 plate with or without 1 μM CHL. Pictures were taken with Nikon D60 digital camera. For the luciferase reporter assays, images were captured with an Amersham imager 600, or Canon 5D Mark IV (33).

### Pre-exposure MIC Assay

Bacillus subtilis cultures were grown overnight in 4 mL LB broth at 37 °C with shaking. Overnight cultures were diluted to an OD_600_ of 0.08 in fresh LB. Cultures were then divided into two groups: one was grown in the presence of CHL (1 µM), while the other served as an unexposed control. Both groups were grown for 2.5 hours (OD_600_ of ctrl = 1) before proceeding to MIC assays. For each experimental condition, cultures were inoculated with 5 × 10^5^ cells per well from pre-exposed and nonexposed (ctrl) cultures and transferred to a 96-well plate. A two-fold serial dilution series of (32 µg/mL to 0.03125 µg/mL) of CHL, lincomycin, tylosin, or phleomycin were added across wells. Plates were incubated in an Agilent BioTek Synergy H1 microplate reader at 30 °C with continuous orbital shaking. OD_600_ was monitored every 30 minutes for 18 hours. All assays were performed with four biological replicates per condition.

### Modified Kirby-Bauer Assays

GYM7 agar plates were prepared one day before the experiment. Agar plates were dried under the hood for 25 min before use. Overnight LB cultures were diluted to 0.08 OD_600_, then grown to 1.0. 100 μL of respective *B. subtilis* cultures (OD_600_=1.0) were spread on the plate using glass beads and then dried for 5 min with the petri dish lid open. Paper discs (6 mm) were then placed on the GYM7 plate, followed by adding 10 μL each antibiotic. Plates were incubated at 30°C for 24 hours.

## Acknowledgements

Thank Jennifer Herman’s lab for advice and strains. Thank you to Jonathan Cortez, Aya Teymur, and Sankirthana Malireddy for contributing to this project. Research reported in this publication was supported by the National Institute of General Medical Sciences of the National Institutes of Health under award number R01GM141700. Sandra LaBonte was support in part by NIH Training Grant T32 GM135115.

---

**Figure S1.**
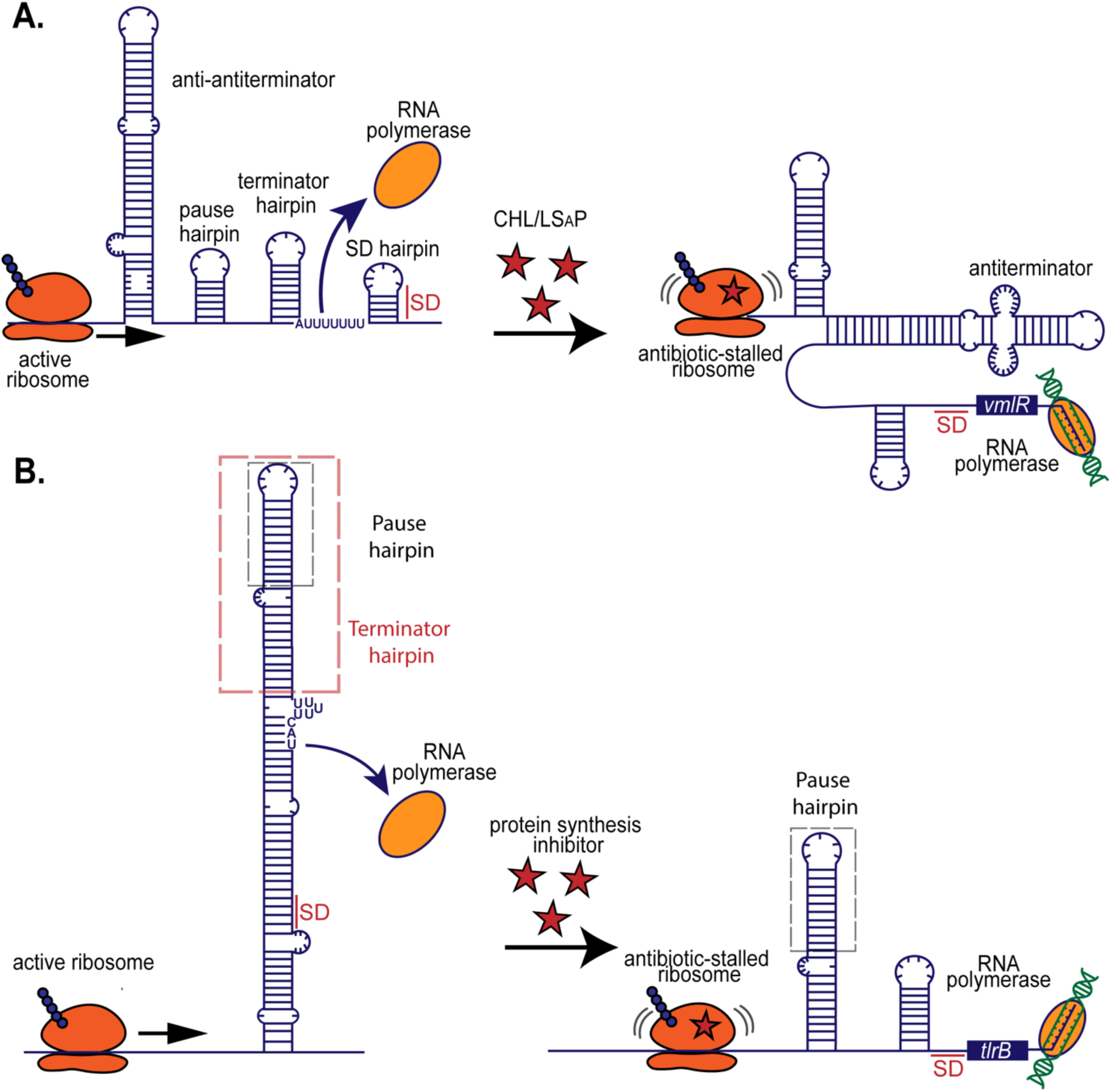
Models for VmlR and TlrB regulation Overview of the leader peptide transcription attenuation regulatory mechanisms for *vmlR* (**A**) and *tlrB* (**B**). **A.** In the absence of drug, the leader peptide nascent transcript forms 4 stem loops: an anti-antiterminator hairpin, a NusG-dependent pause hairpin, a terminator hairpin, and, if there is leaky transcription past the terminator hairpin, a Shine-Dalgarno-sequestering (SD) hairpin that prevents ribosome binding. Expression of *vmlR* is induced in the presence of CHL or L_S_APs. These protein synthesis inhibitors stall the ribosome causing only partial formation of the first stem loop and formation of antiterminator structure. Therefore, transcription proceeds to the VmlR ORF (15). **B.** The regulation of *trlB* is more complex than *bmrCD* and *vmlR*. In the absence of antibiotic, NusG-dependent pausing allows for initiation of *tlrB* uORF translation. The ribosome interferes with the formation of a terminator hairpin when reaching the uORF stop codon, but then releases, and termination occurs. Transcripts evading termination are suppressed due to the stem loop sequestering the *tlrB* SD sequence. In the presence of tylosin or other translation inhibitors, however, the ribosome stalls on the uORF where the terminator hairpin would begin to form. Transcription termination is therefore suppressed. Additionally, the SD sequence is no longer trapped within a stem loop structure, and the ribosome can bind the transcript. Therefore, *tlrB* is transcribed and subsequently translated (12).

**Figure S2.**
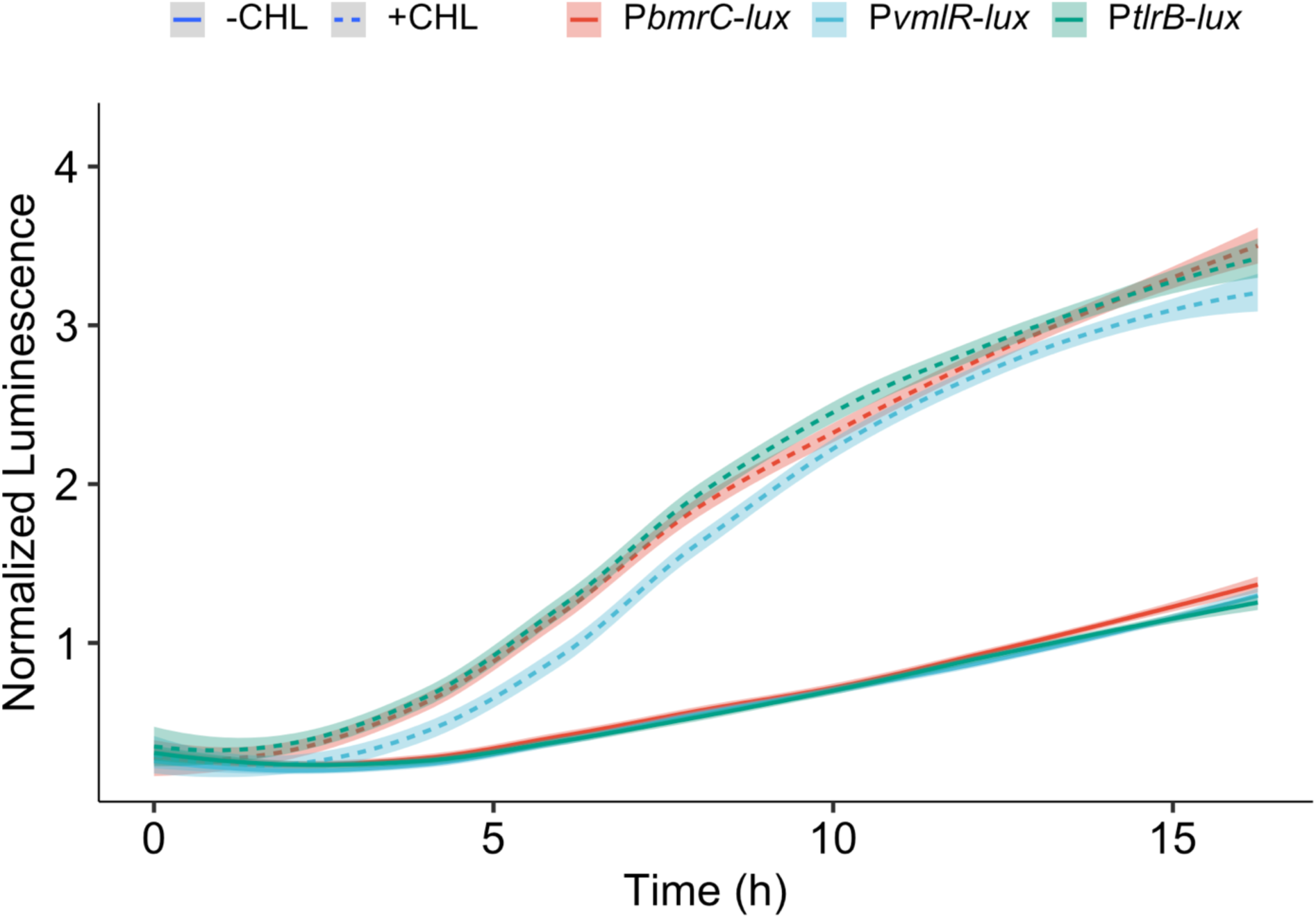
Transcriptional activity of 5’uORF-luciferase fusions P*bmrC-lux,* P*vmlR-lux,* and P*tlrB-lux* in relative light units (RLU) exposed to 1 μM CHL (dashed) compared to non-exposed (solid) grown over 16 hours. The curves were normalized to a luciferase construct lacking a promoter.

**Figure S3.**
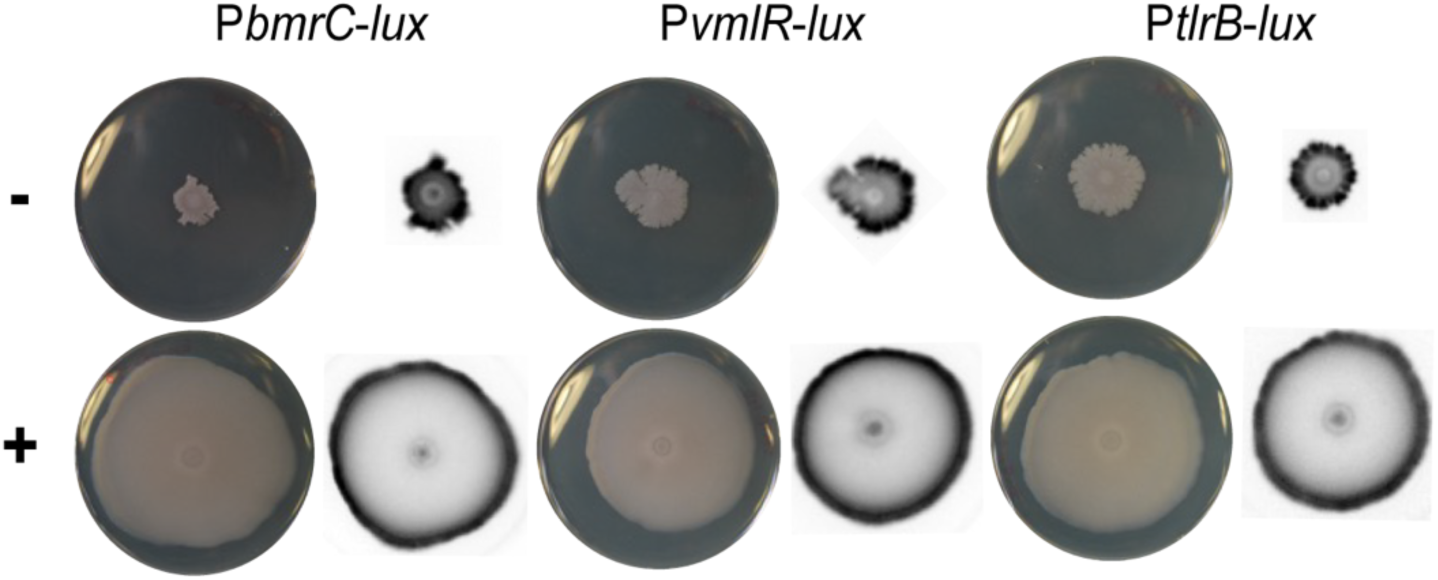
Transcriptional activity of resistance genes at 48 hours of growth (**Note:** The chemiluminescence images are inverted. The luminescence signal appears gray to black with increasing intensity) P*bmrC-lux,* P*vmlR-lux,* and *PtlrB-lux* spotted on agar plates with 1 μM CHL (+) or without CHL (-). Pictures were taken with phase contrast (left) and chemiluminescence (right) at 48 hours of growth.

**Figure S4.**
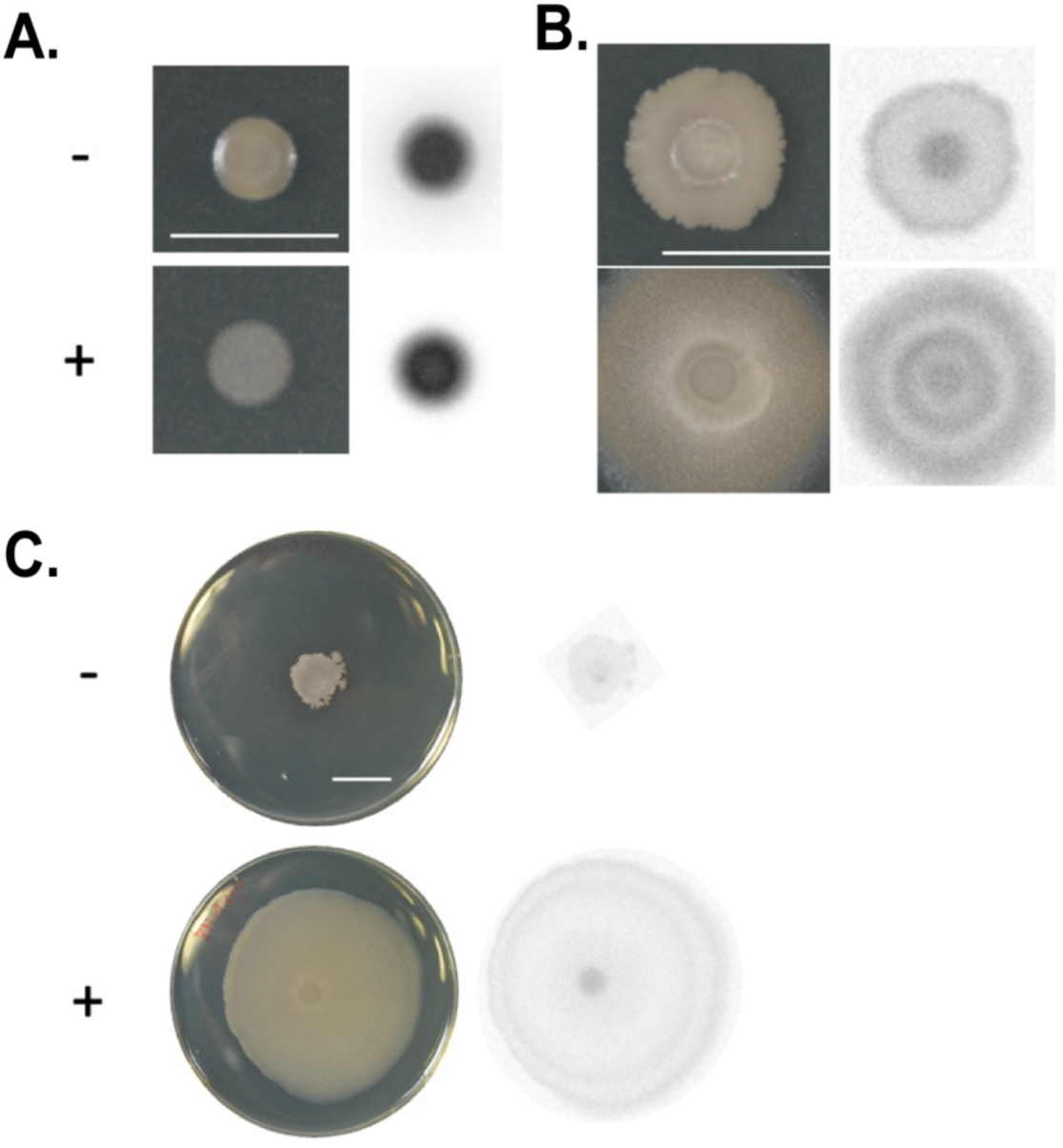
Transcriptional activity of *ytbDE* over 24 hours of growth (**Note:** The chemiluminescence images are inverted. The luminescence signal appears gray to black with increasing intensity) P*ytbDE-lux* was spotted on agar plates with 1 μM CHL (+) or without CHL (-). Pictures were taken with phase contrast (left) and chemiluminescence (right) at **A.** 6 hours, **B.** 24 hours of growth.

**Figure S5.**
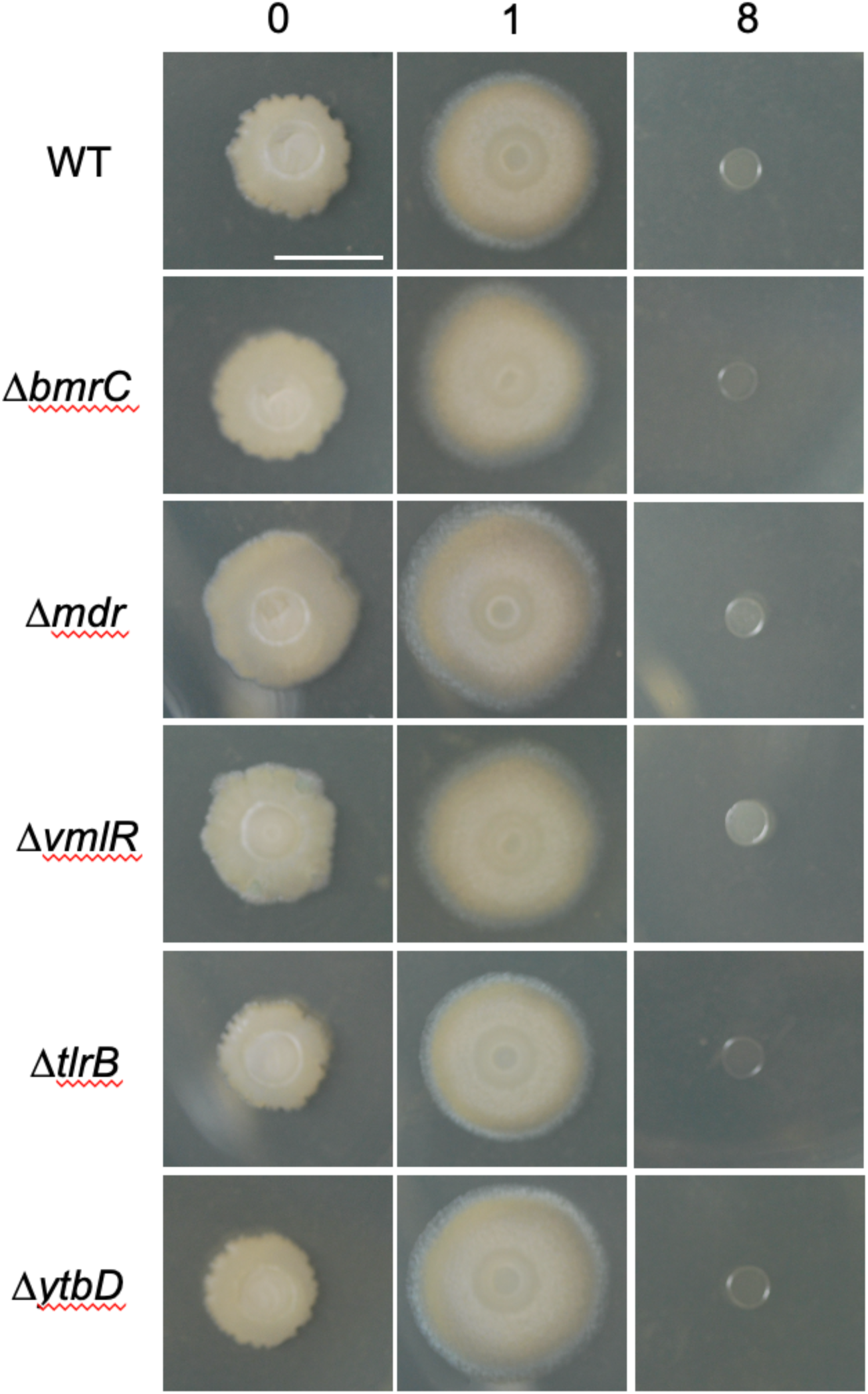
Single gene deletions of each resistance gene do not provide resistance or change colony morphology in response to CHL exposure. Wild type (WT) and each deletion strain were plated on the GYM7 plate with no CHL, subinhibitory CHL (1 μM) and inhibitory CHL (8 μM). Pictures were taken at 24 h.

**Figure S6.**
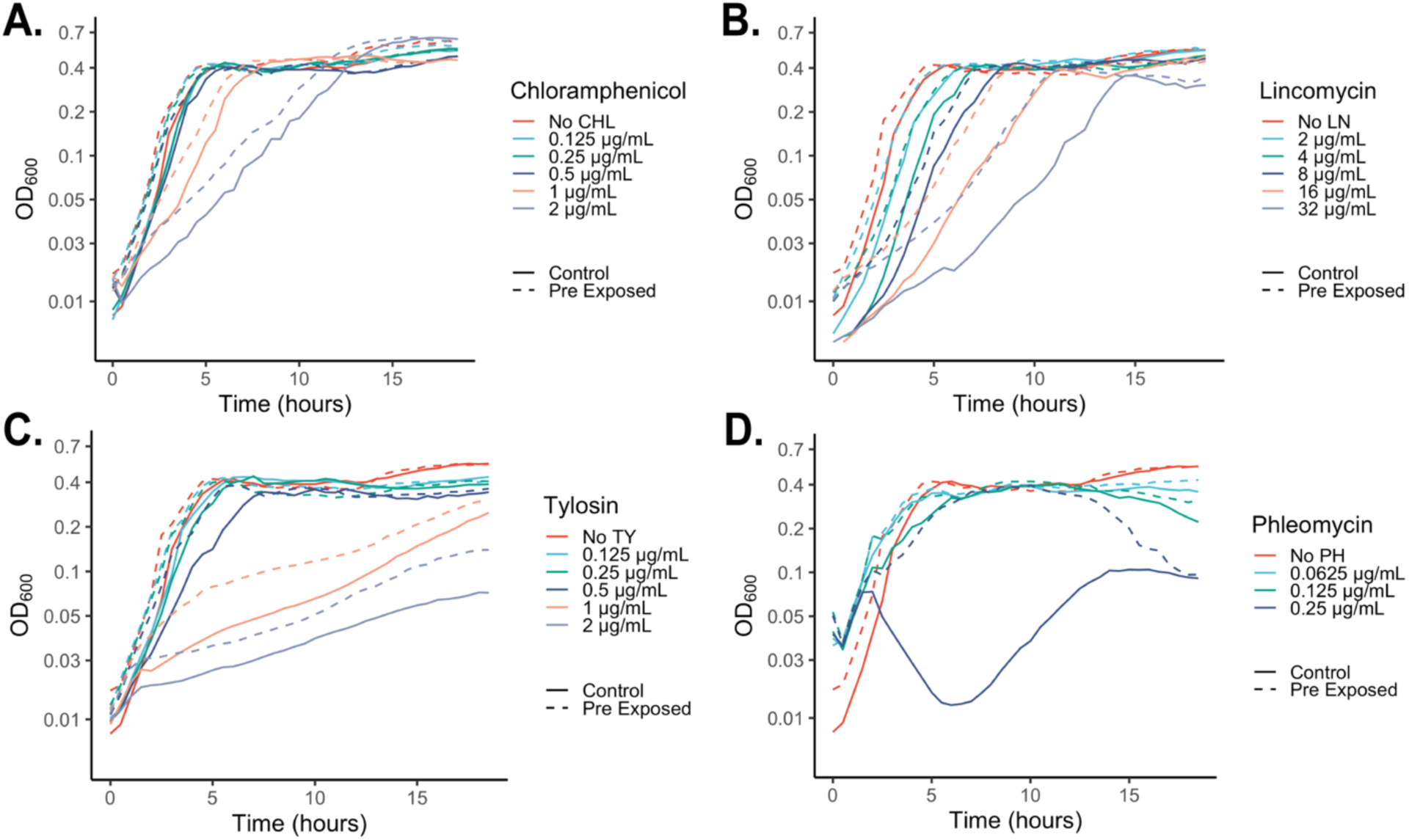
Growth comparison of cultures pre-exposed to CHL (dashed) to cultures grown in isolation (solid) at a range of Chloramphenicol (**A**), Lincomycin (**B**), Tylosin (**C**), and Phleomycin (**D**) concentrations over 18 hours of growth. OD_600_ values were taken every 30 minutes. Each line is an average of 4 replicates.

**Figure S7.**
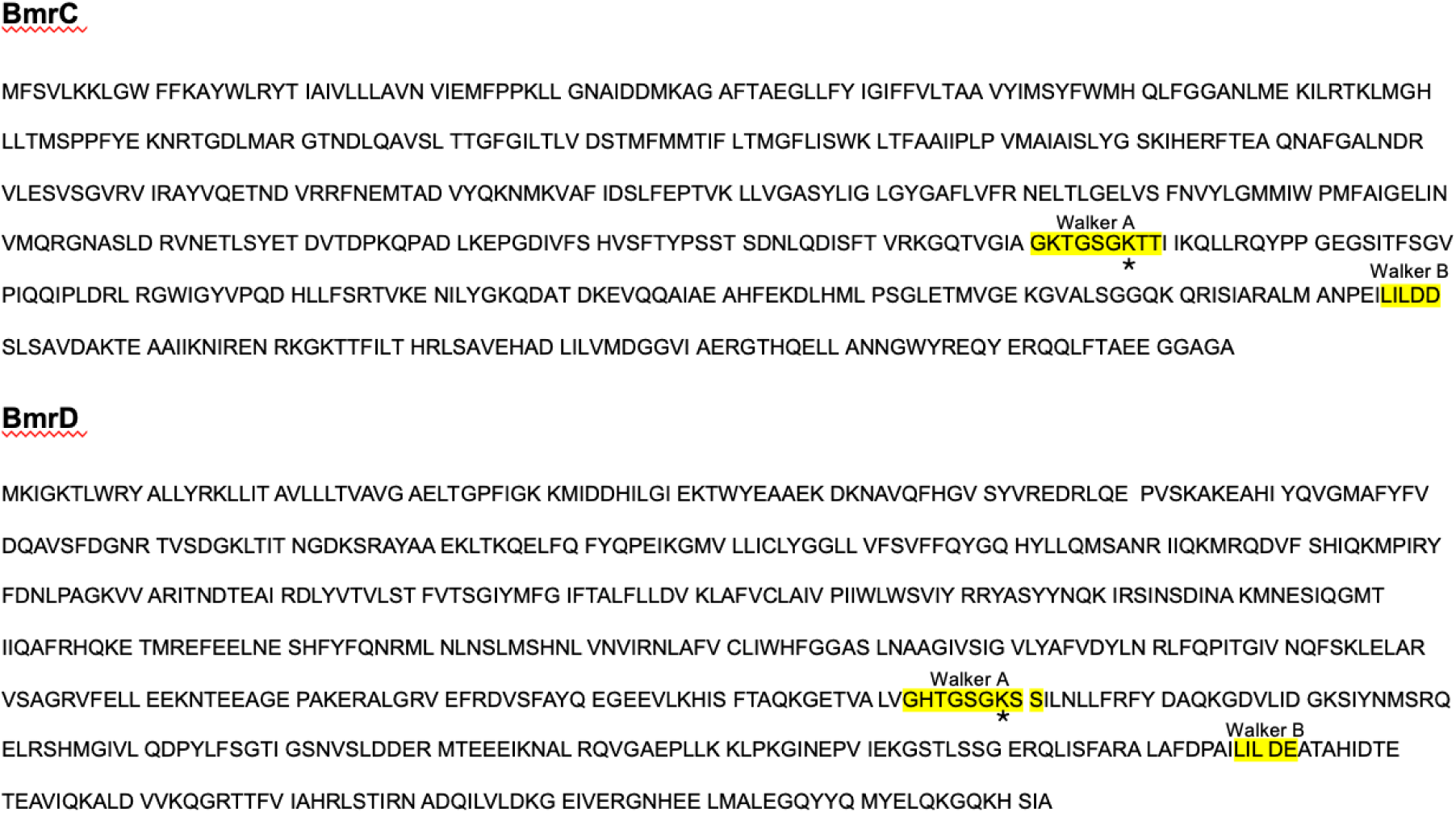
Walker A and B motifs of BmrCD. BmrC and BmrD protein sequences with the Walker motifs highlighted in yellow. The sites mutated in this study (BmrC K377A; BmrD K469A) are indicated below the residue with an asterisk.

**Figure S8.**
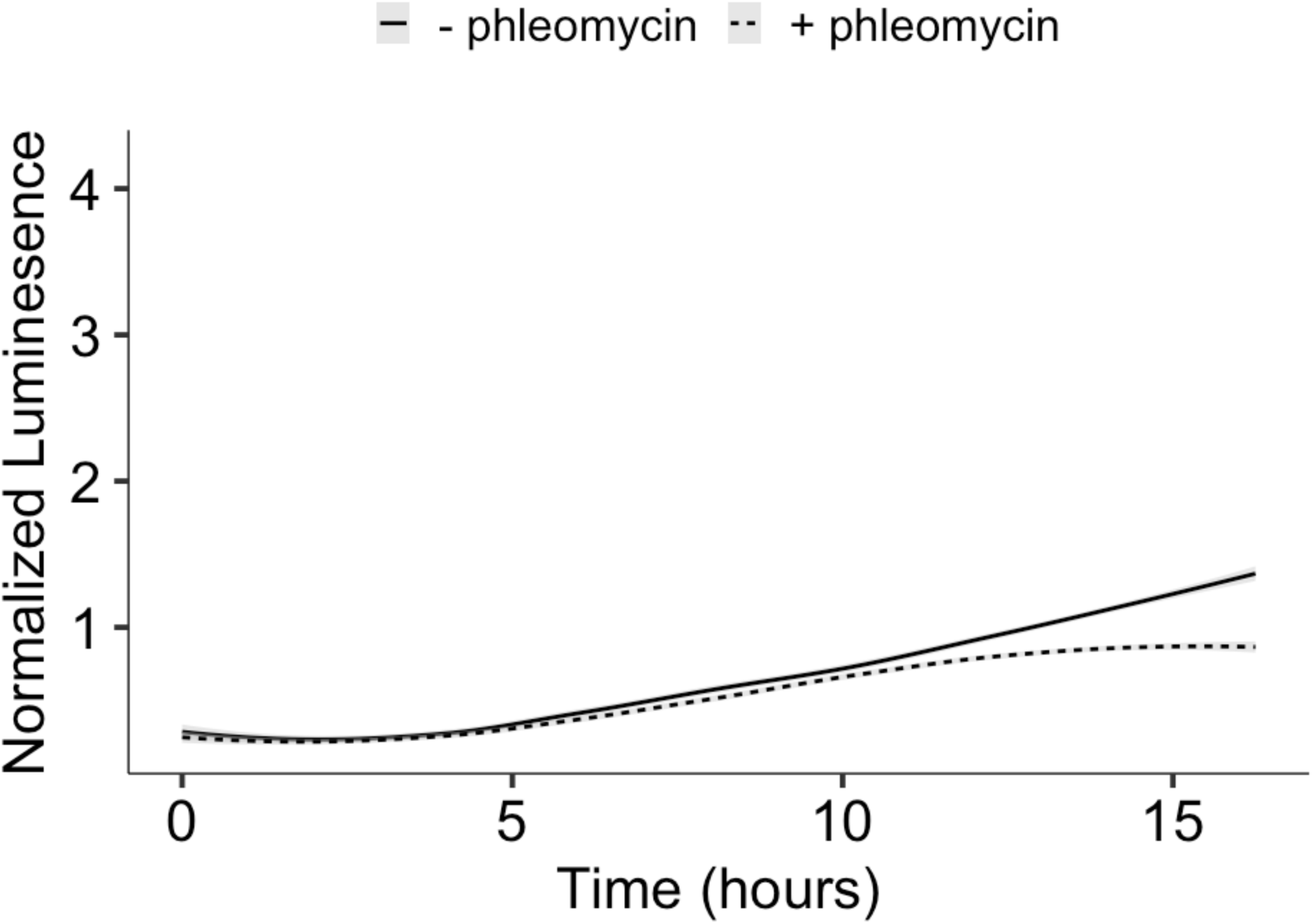
BmrCD expression is not triggered by phleomycin exposure. We monitored luminescence over 16 hours of P*bmrC-lux* grown with (dashed) and without (solid) 0.125 μg/mL phleomycin. Curves are normalized to a control luciferase construct lacking a promoter. Each line is average of 4 replicates.

**Table S1.** Antibiotics tested for resistance via *bmrCD and ytbDE*.

| **Antibiotic** | **Class** |
| --- | --- |
| Apramycin | Aminoglycoside |
| Kanamycin | Aminoglycoside |
| Hygromycin B* | Aminoglycoside |
| Erythromycin | Macrolide |
| Tylosin* | Macrolide |
| Spectinomycin | Aminocyclitol |
| Ampicillin* | Penicillin |
| Carbenicillin* | Penicillin |
| Lincomycin | Lincosamide |
| Tetracycline | Tetracycline |
| Chloramphenicol | Amphenicol |
| Rifamycin** | Ansamycin |
| Phleomycin** | Glycopeptide |

\*Only tested for *ytbDE*
\*\*Only tested for *bmrCD*

**Table S2.**
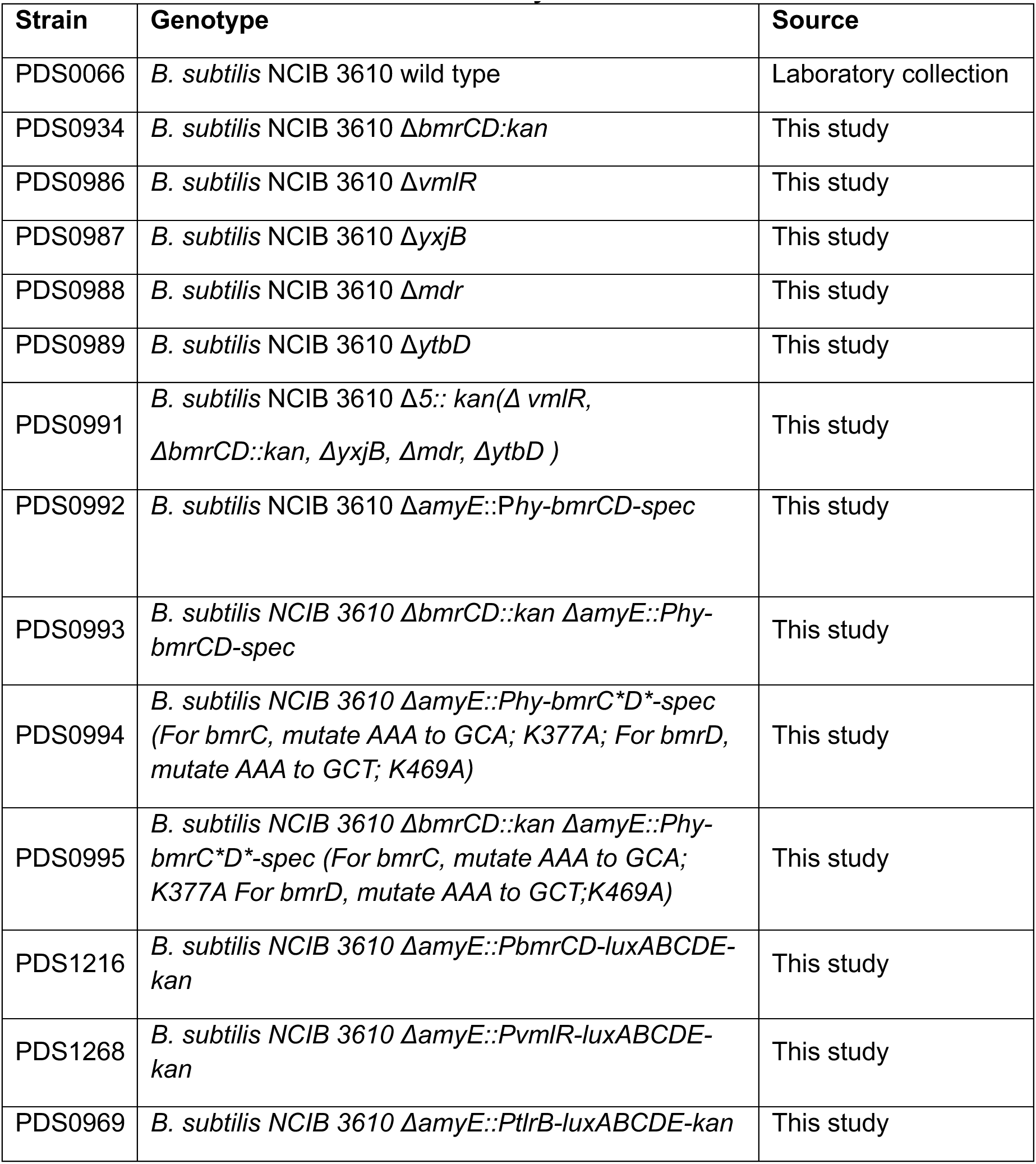
Bacterial strains used in this study.

**Table S3.**
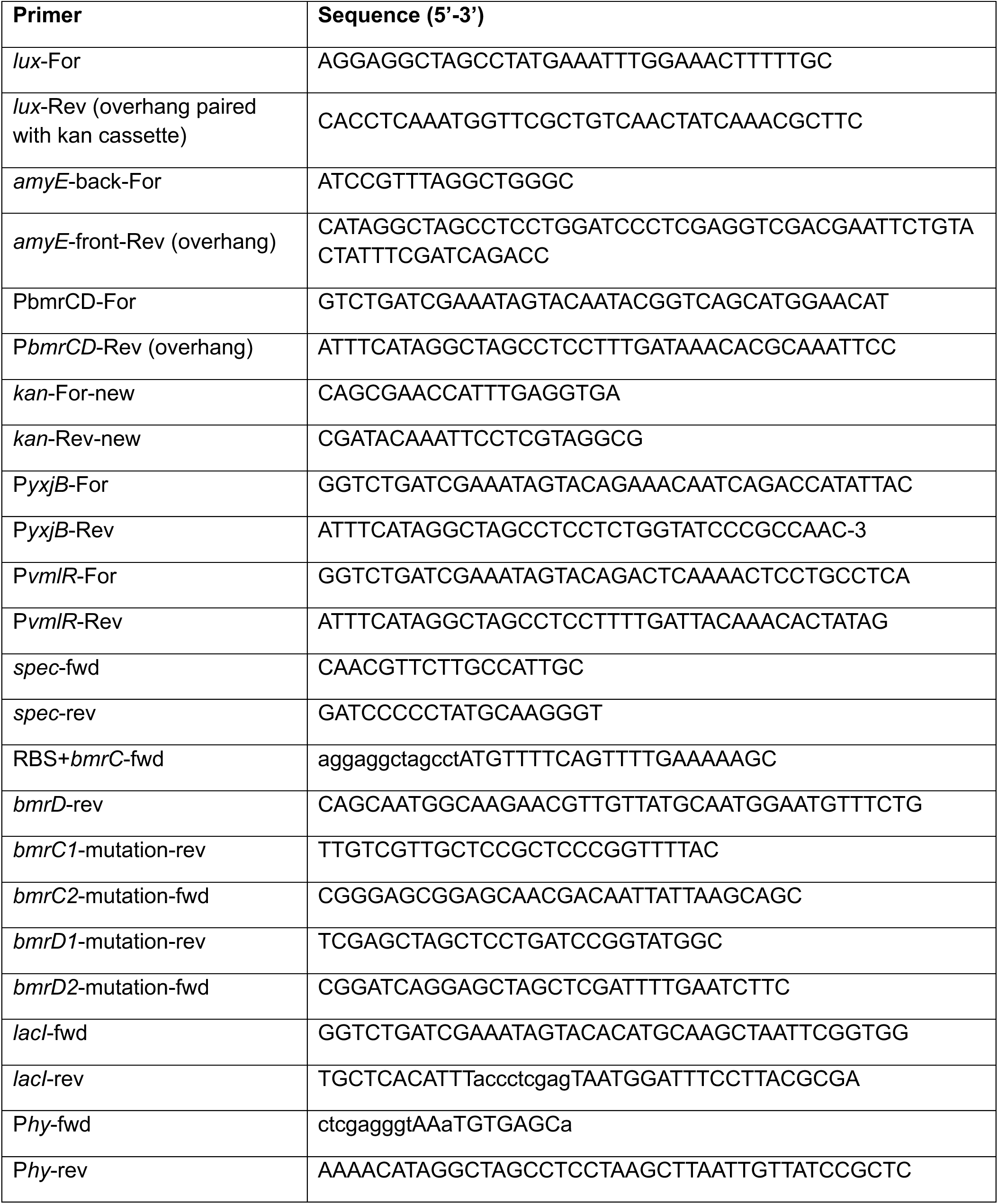
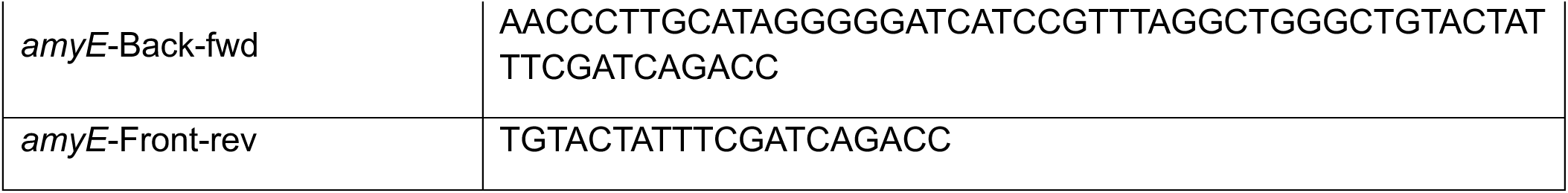
Primers used in this study.

